# Changes in cell morphology and function induced by *NRAS* Q61R mutation in lymphatic endothelial cells

**DOI:** 10.1101/2023.07.14.549027

**Authors:** Shiho Yasue, Michio Ozeki, Akifumi Nozawa, Saori Endo, Hidenori Ohnishi

## Abstract

Recently, a low-level somatic mutation in *NRA*S gene (c.182 A > G, Q61R) was identified in the specimens of patients with kaposiform lymphangiomatosis (KLA). However, it is unknown how these low-frequency mutated cells can affect the characterization and surrounding environment of their lesions. To understand the pathogenesis and association of these gene abnormalities, we established *NRAS*^Q61R^ mutated lymphatic endothelial cells (LECs) transfected with lentivirus vector and undertook morphological and functional characterization, protein expression profiling, and metabolome analysis. *NRAS*^Q61R^ human dermal LECs showed poor tube formation and high cell proliferation and migration ability with increasing ratios of mutated cells. Analysis of signaling pathways showed inactivation of the PIK3/AKT/mTOR pathway and hyperactivation of the RAS/MAPK/ERK pathway, which was improved by MEK inhibitor treatment. This study may show the theoretical circumstances *in vitro* induced by *NRAS*^Q61R^-mutated cells in the affected lesions of KLA patients.

## Introduction

Generalized lymphatic anomaly (GLA) and kaposiform lymphangiomatosis (KLA), known as complex lymphatic anomalies (CLAs), are rare multiorgan diseases that cause pleural effusion, pericardial effusion, ascites, and osteolysis due to the proliferation of abnormal lymphatic tissue^1^. In the updated International Society for the Study of Vascular Anomalies 2018 classification, GLA is classified as lymphatic malformations (LMs)^2^. KLA is a subtype of GLA characterized by spindle-shaped lymphatic endothelial cells (LECs) and presents with more serious symptoms (abnormal coagulation, thrombocytopenia, and hemorrhagic pericardial and pleural effusion) and poorer outcomes than GLA patients^3^. However, both the etiology and pathogenesis of KLA have yet to be clarified.

Although genetic analysis has identified low-level somatic mutations in the specimens of patients with vascular anomalies, little has been reported on the associated genetic abnormalities in KLA. Nonetheless, mutations in genes encoding components of the PIK3/AKT/mTOR and RAS/MAPK/ERK pathways have recently been identified in the affected lesions of CLA patients. A somatic mutation in *NRA*S (c.182 A > G, Q61R) was detected in the lesions in 10/11 cases of KLA^4^. Our group also detected an *NRAS* mutation in the cell-free DNA of the plasma and pleural effusion of KLA patients^5^. The frequencies of the mutated alleles in the affected cells in these reports were very low (lower than 30% or only in a few cells). However, it is still unknown as to how the low-frequency mutated cells can affect the characteristics and the surrounding environment of their lesions.

To understand the pathogenesis and association of these gene abnormalities, animal models and patient-derived cells have been studied^6–8^. Studies in mouse models have provided important findings regarding the mechanisms of *PIK3CA* mutations affecting the phenotype of LECs and vessel overgrowth. LM patient-derived LECs with mutant *PIK3CA* showed increased proliferation and resistance to stimuli on cell death^9^. Mutated *PIK3CA*-expressing murine LECs also showed increased migration and sprouting^10^. Thus, the development of LMs is known to be triggered by a somatic activating *PIK3CA* mutation in LECs, leading to cell autonomous proliferation and migration caused by activation of PIK3/AKT signaling. However, most LM cells in affected lesions that show abnormal morphology, both grossly and microscopically, do not have any genetic abnormalities. Therefore, little is known about how the very small proportion of mutated LECs can affect other non-mutated cells or surrounding environments.

In this study, we established *NRAS*^Q61R^-mutated LECs transfected with lentivirus vector and performed morphological and functional characterizations, protein expression profiling, and metabolome analysis. To elucidate the influence of the mutated LECs on wild type LECs, we studied the effects of mixing these cell populations. Furthermore, we examined the response of these cells to mTOR and MAPK kinase (MEK) inhibitors.

## Materials and Methods

### Cell and vector preparation

Juvenile foreskin-derived human dermal lymphatic endothelial cells (HDLECs) were purchased from PromoCell (Heidelberg, Germany). Cells were cultured in Endothelial Cell Growth Basal Medium-2 (EBM2) (Lonza, Basel, Switzerland) with 10% heat inactivated fetal bovine serum, penicillin (100 units/mL), and streptomycin (100 μg/mL) and propagated at 37°C under 5% CO₂.

cDNA of *NRAS*^Q61R^ (accession number: NM_002524.5) with a 3’ terminal 6 × His-tag was chemically synthesized and cloned into the pFastBac1 vector (Invitrogen, Carlsbad, California, USA). cDNA of *NRAS*^Q61R^ was cloned into the pLVSIN-EF1α-AcGFP-C1 vector (Takara Bio Inc., Otsu, Japan).

### Gene transfection of HDLEC cell line using lentiviruses

Using the Lenti-X 293 T cell line (Takara Bio Inc., Otsu, Japan) and the pLVSIN-EF1α-AcGFP-C1 vector, we transfected *NRAS* ^Q61R^ vectors into HDLECs. The Lenti-X 293 T cell line was spread on 10-cm plates at a density 5.0 × 10^6^ cells per plate and cultured in Dulbecco’s Modified Eagle Medium (DMEM) for 24 h. The pLVSIN-EF1α-AcGFP-C1 vector (5.5 μg) was added to 7 μL Lentiviral Mix High Titer Packaging Mix (Takara Bio Inc., Otsu, Japan), 1500 μL serum-free DMEM, and 45 μL Trans IT-293 Transfection Reagent (Takara Bio Inc., Otsu, Japan). After 15 min, this mixture was added to Lenti-X 293 T cells and incubated at 37°C. After 24 h, the medium was exchanged with DMEM. After 48 h, the culture medium was collected and passed through a 0.45-μm filter (this solution contained recombinant lentiviruses). HDLECs were spread onto six-well plates at a density of 2.0 × 10^5^ cells per well. The recombinant lentiviral solution was diluted with EBM2 to give solutions with initial concentrations of 1:4 to 1:10. The medium was discarded from each well, and 2 ml of the recombinant lentiviral solutions were added to each well. Polybrene was added to the medium to a final concentration of 4 μg/ml. To select stably GFP-only or C-terminal GFP-fused *NRAS*^Q61R^ cells, repeated exchange of the medium and the addition of puromycin (final concentrations of 1.5 μg/m and 0.5 μg/ml, respectively) were performed every 72 h. *NRAS*^Q61R^ expression in these cells was checked by polymerase chain reaction (PCR). *GFP* gene was transfected into HDLECs by the same method. These transfected cells and *NRAS* wild type fluorescent cells were used in the scratch assay.

### Analysis of NRAS mutational status

Genomic DNA was extracted from cell pellets using the Sepa Gene Kit (EIDIA Co., Ltd., Tokyo, Japan) in accordance with the manufacturer’s instructions. DNA extraction was performed using a single extraction method. After extraction, DNA was stored at 4°C until use.

The following primers were used for SNP analysis: *NRAS*-forward 5’-CAGGTGGTGTTGGGAAAAGC-3’; *NRAS*-reverse 5’-CTCGCTTAATCTGCTCCCTGT-3’. The sequences were analyzed using a BigDye Terminator v1.1 Cycle Sequencing Kit and an Applied Biosystems 3130xl Genetic Analyzer (Applied Biosystems, MA, USA).

### Tube formation assay

Cells were plated on thin layers of Geltrex (Thermo Fisher Scientific, MA, USA) at a density of 1 × 10^4^ cells/well in a 96-well plate, cultured in EBM2, and incubated at 37°C for 20 h. Images were captured automatically every 5 min using time-lapse video microscopy (WSL-1800-B; ATTO Corp., Tokyo, Japan). After 20 h, microscopic images were taken and analyzed by Wimasis Image Analysis (WimTube; Wimasis GmbH Munich, Germany). Tube formation was observed by microscopy (BZ-9000; KEYENCE, Osaka, Japan). To assess the association between the proportion of mutated cells and the function of tube formation, the differences in tube formation function were examined in *NRAS* wild type HDLECs (*NRAS*^WT^) (*NRAS*^WT^ 100% and *NRAS*^Q61R^ 0%), *NRAS*^Q61R^ HDLECs (*NRAS*^WT^ 0% and *NRAS*^Q61R^ 100%), and mixed HDLECs (*NRAS*^WT^ 95% and *NRAS*^Q61R^ 5%, and *NRAS*^WT^ 25% and *NRAS*^Q61R^ 75%). To analyze the function of tube formation, covered area, total nets, and tube length standard deviation in each image was counted and average values within groups were compared. “Covered area” means the area occupied by cellular components, and “total nets” means the number of nets (a structure in which two or more loops are connected). At least three fields per well were examined, and each experimental condition was tested in triplicate. In addition, 30 ng/ml mTOR inhibitor (rapamycin (sirolimus); Sigma–Aldrich, St. Louis, MO, USA) or 30 ng/ml MAPK kinase (MEK) inhibitor (trametinib; ChemScene, Monmouth Junction, NJ, USA) was added to these cells for examination of drug inhibition.

### Scratch assay

*NRAS*^WT^ and *NRAS*^Q61R^ HDLECs were cultured to 90% confluency in 24-well plates. Cell monolayers were scraped with a 200-μl pipette tip to make a wound, and then the cells were cultured in EBM2. Images were captured automatically every 5 min using time-lapse video microscopy (WSL-1800-B; ATTO Corp., Tokyo, Japan). Quantification of wound areas was performed using image analysis software (Image J) at 0, 6, 12, 18, and 24 h.

### Cell proliferation assay

A Cell Counting Kit-8 assay (CCK-8; Dojindo Laboratories, Kumamoto, Japan) was used to assess the effects of mTOR inhibitor sirolimus or MEK inhibitor trametinib on cell proliferation, in accordance with the manufacturer’s protocol. We seeded 1 × 10^4^ cells/well in 96-well plates and incubated them with sirolimus (40 ng/ml), trametinib (20 ng/ml), or vehicle control (dimethyl sulfoxide) for 48 h. To assess the association between the proportion of mutated cells and cell proliferation, the differences were examined in *NRAS*^WT^, *NRAS*^Q61R^, and mixed HDLECs. We then added 10 µl CCK8 solution to each well with 100 µl medium and incubated the cells for 3 h. After incubation, absorbance was measured at 450 nm using a plate reader (model 680; Bio-Rad Laboratories, Inc., Hercules, CA, USA).

### Analysis of signaling pathways by western blotting and multiplex protein assay

Cells were lysed in cold lysis buffer (CytoBuster Protein Extraction Reagent, Novagen Inc., Madison, WI, USA) in the presence of a protease inhibitor cocktail (Complete Mini, RocheDiagnostics, Mannheim, Germany). Protein samples were separated by sodium dodecyl sulfate polyacrylamide gel electrophoresis and transferred to polyvinylidene fluoride membranes. The membranes were incubated with 5% skimmed milk solution and probed with the indicated primary antibodies at 4°C overnight. Membranes were then washed and incubated with secondary antibodies for 1 h at room temperature. Protein bands were visualized using ECL Prime Western Blotting Detection Regent (GE Healthcare; Amersham, UK) and a light-capture cooled CCD camera system (ATTO Corp., Tokyo, Japan). Monoclonal antibodies against AKT (2920S) and polyclonal antibodies against phospho-AKT (Ser473) (p-AKT [Ser473]; 9271S) were purchased from Cell Signaling Technology (Boston, MA, USA). Polyclonal antibodies against extracellular signal-regulated kinase 1/2 (ERK1/2; ab196883) and monoclonal antibodies against phospho-ERK1/2 (p-ERK1/2; ab201015) were purchased from Abcam. Protein expression was normalized to GAPDH (sc-2577; Santa Cruz Biotechnology, Santa Cruz, CA, USA)

11-Plex Akt/mTOR Total Protein Magnetic Bead Kit (cat #: 48-612MAG; MILLIPLEX), Akt/mTOR Phosphoprotein 11-plex Magnetic Bead (cat #: 48-611-MAG; MILLIPLEX), and Phospho/Total ERK 2-Plex Magnetic Bead Kit (EMD Millipore Corporation, Billerica, MA, U.S.A.) were used to assess the levels of these proteins. Cells were positioned in six-well plates and cultured until 90% confluency. To assess the association between the proportion of mutated cells and the levels of these proteins, the differences were examined in *NRAS*^WT^, *NRAS*^Q61R^, and mixed HDLECs in triplicate at one time using a single plate. The procedure was performed in accordance with the manufacturer’s assay protocols (EMD Millipore, Billerica, MA, USA). A Luminex 200 machine and MILLIPLEX Analyst software were used for data analysis.

### Suspension array

The concentrations of 16 angiogenic and lymphangiogenic factors in the culture medium of *NRAS*^WT^ and *NRAS*^Q61R^ HDLECs were evaluated using a commercial kit from Luminex (Supplemental Table 1). Culture medium without cells was used as the control. The Human Angiogenesis/Growth Factor Magnetic Bead Panel (MILLIPLEX MAP, HAGP1MAG-12K) was performed in duplicate in accordance with the manufacturer’s protocol (EMD Millipore, Billerica, MA, USA).

### Metabolite extraction

Culture medium was aspirated from the dish and the cells were washed twice with 5% mannitol solution (10 ml and 2 ml for the first and second washes, respectively). The cells were then treated with 800 µl methanol and incubated at room temperature for 30 sec to suppress enzymatic activity. Next, 550 µl Milli-Q water containing internal standards (H3304-1002, Human Metabolome Technologies, Inc. (HMT), Tsuruoka, Yamagata, Japan) was added to the cell extract, followed by further incubation at room temperature for 30 sec. The cell extract was then centrifuged at 2,300 ×*g* for 5 min at 4°C, after which 700 µl of the supernatant was centrifugally filtered through a Millipore 5-kDa cutoff filter (UltrafreeMC-PLHCC, HMT) at 9,100 ×*g* for 120 min at 4°C to remove macromolecules. Subsequently, the filtrate was evaporated to dryness under vacuum and reconstituted in 50 µl Milli-Q water for metabolome analysis at HMT.

### Metabolome analysis (C-SCOPE)

Metabolome analysis was conducted in accordance with HMT’s *C-SCOPE* package, using capillary electrophoresis time-of-flight mass spectrometry (CE-TOFMS) for cation analysis and CE-tandem mass spectrometry (CE-MS/MS) for anion analysis on the basis of the methods described previously^11, 12^. Briefly, CE-TOFMS and CE-MS/MS analyses were performed using an Agilent CE capillary electrophoresis system equipped with an Agilent 6210 time-of-flight mass spectrometer (Agilent Technologies, Inc., Santa Clara, CA, USA) and Agilent 6460 Triple Quadrupole LC/MS (Agilent Technologies), respectively. The systems were controlled by Agilent G2201AA ChemStation software version B.03.01 for CE (Agilent Technologies) and connected by a fused silica capillary (50 μm *i.d.* × 80 cm total length) with commercial electrophoresis buffer (H3301-1001 and I3302-1023 for cation and anion analyses, respectively, at HMT) as the electrolyte. The time-of-flight mass spectrometer was scanned from *m/z* 50 to 1,000^11^ and the triple quadrupole mass spectrometer was used to detect compounds in dynamic MRM mode. Peaks were extracted using MasterHands, automatic integration software (Keio University, Tsuruoka, Yamagata, Japan)^13^ and MassHunter Quantitative Analysis B.04.00 (Agilent Technologies) to obtain peak information including *m/z*, peak area and migration time (MT). Signal peaks were annotated in accordance with HMT’s metabolite database based on their *m*/*z* values and MTs. The peak area of each metabolite was normalized to internal standards, and metabolite concentration was evaluated by standard curves with three-point calibrations using each standard compound. Hierarchical cluster analysis and principal component analysis^14^ were performed by HMT’s proprietary MATLAB and R programs, respectively. Detected metabolites were plotted on metabolic pathway maps using VANTED software^15^.

### Statistical analysis

Welch’s t-test was used to compare two groups, and one-way ANOVA was used to compare three or more groups. For comparisons of three or more groups, p-values were calculated by Tukey’s multiple comparison with the 0% group or control group as the reference. P-values were considered significant at <0.05. Data were analyzed using EZR (Easy R) software.

## Results

### *NRAS*^Q61R^ HDLEC characteristics

Gene transfection of *NRAS*^Q61R^ was confirmed by PCR. *NRAS*^Q61R^ HDLECs showed larger and irregular-shaped morphology compared with *NRAS*^WT^ HDLECs microscopically (Fig.1 a–c).

**Figure 1.**
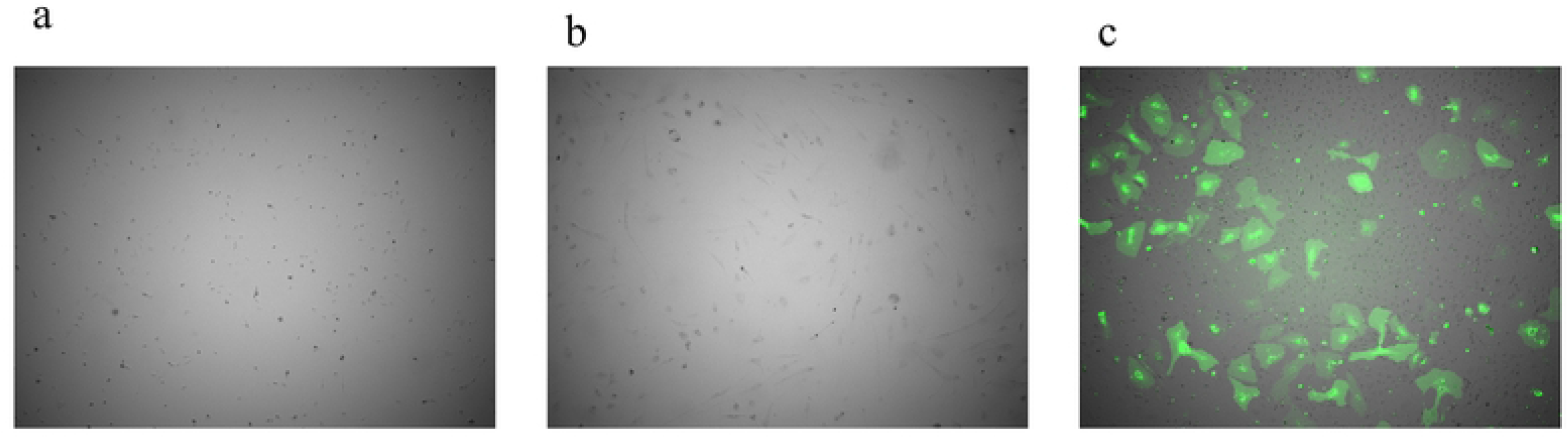
Morphology of human dermal lymphatic endothelial cells (HDLECs) expressing *NRAS*^WT^ and *NRAS*^Q61R^. a: *NRAS*^WT^ HDLECs showed uniform and short-spindle shaped morphology. b: *NRAS^Q61R^* HDLECs showed relatively large and irregular morphology. c: *NRAS*^WT^ and *NRAS^Q61R^* HDLECs were mixed and cultured. *NRAS^Q61R^* HDLECs (fluorescent cells) had larger and more irregular shapes than *NRAS*^WT^ HDLECs.

### Tube formation assay

The tube formation assay revealed that *NRAS*^WT^ HDLECs made well-regulated lumen structures and tube formation, but the sheer-like proliferation patterns and formation of capillary-like structures in *NRAS*^Q61R^ HDLECs were markedly less than those of *NRAS*^WT^ (Figure 2A). In the stepwise examination of *NRAS*^Q61R^ HDLEC mixtures, the covered area, total nets, and tube length standard deviation of each cell line were analyzed. The greater the number of mutated cells, the larger the covered area and the lower the number of total nets (p<0.01) (Fig. 2B). The mean values of the covered area occupying cellular components of the 25% and 100% *NRAS*^Q61R^ HDLEC mixtures were significantly higher than those of the 0% group (p=0.030 and p<0.001, respectively) (Fig. 2B-a). The number of nets of 0% *NRAS*^Q61R^ was significantly lower than that of 5%, 25%, and 100% *NRAS*^Q61R^ (p=0.034, p<0.001, and p<0.001, respectively) (Fig. 2B-b). The standard deviation of tube lengths showed no difference in 0%, 5%, and 25% *NRAS*^Q61R^, but that of 100% *NRAS*^Q61R^ was significantly longer than those of 0%, 5%, and 25% *NRAS*^Q61R^ (p=0.004) (Fig. 2B-c).

**Figure 2.**
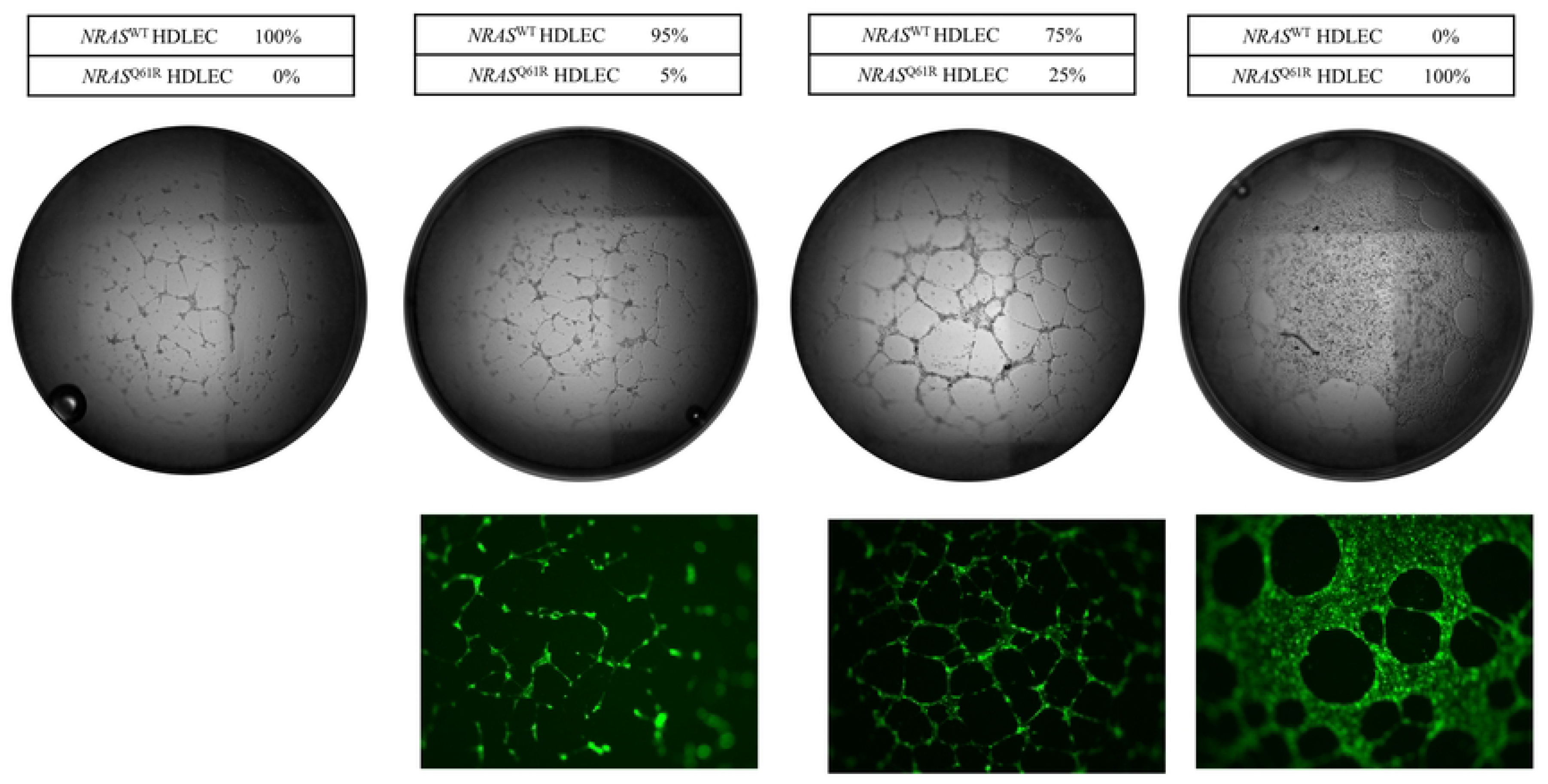
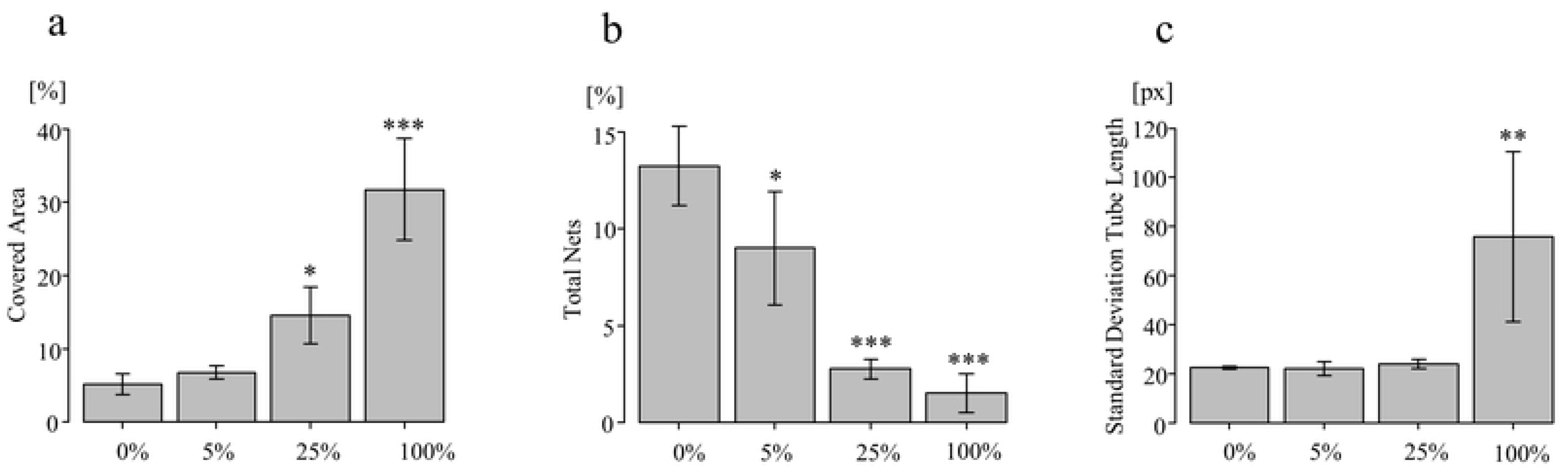
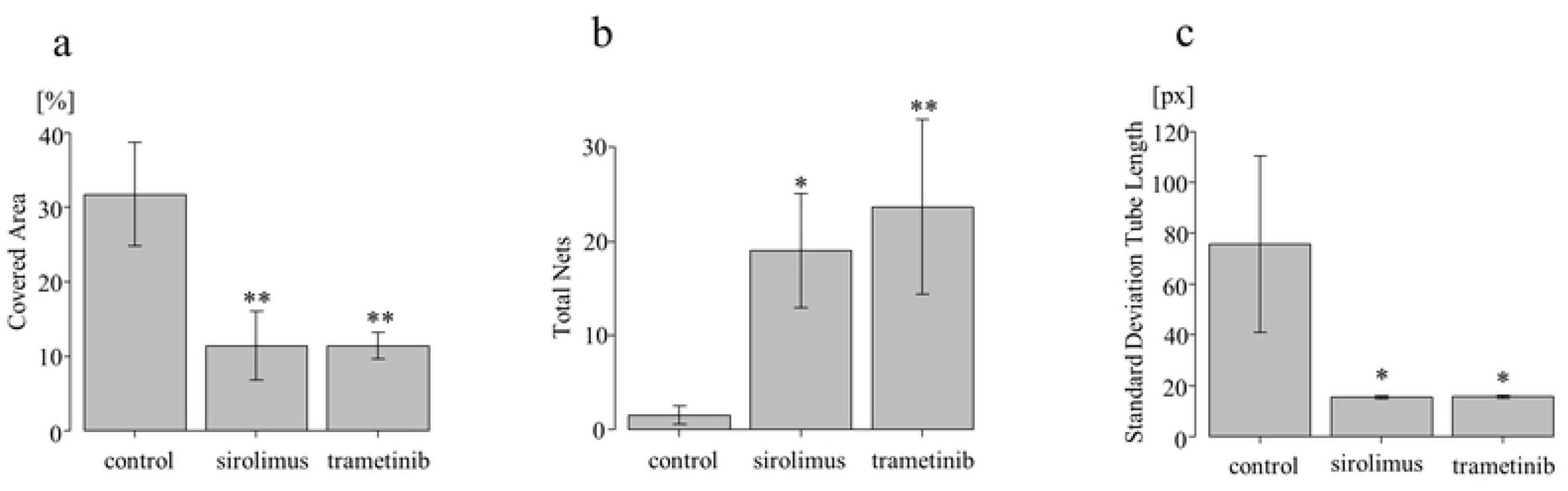
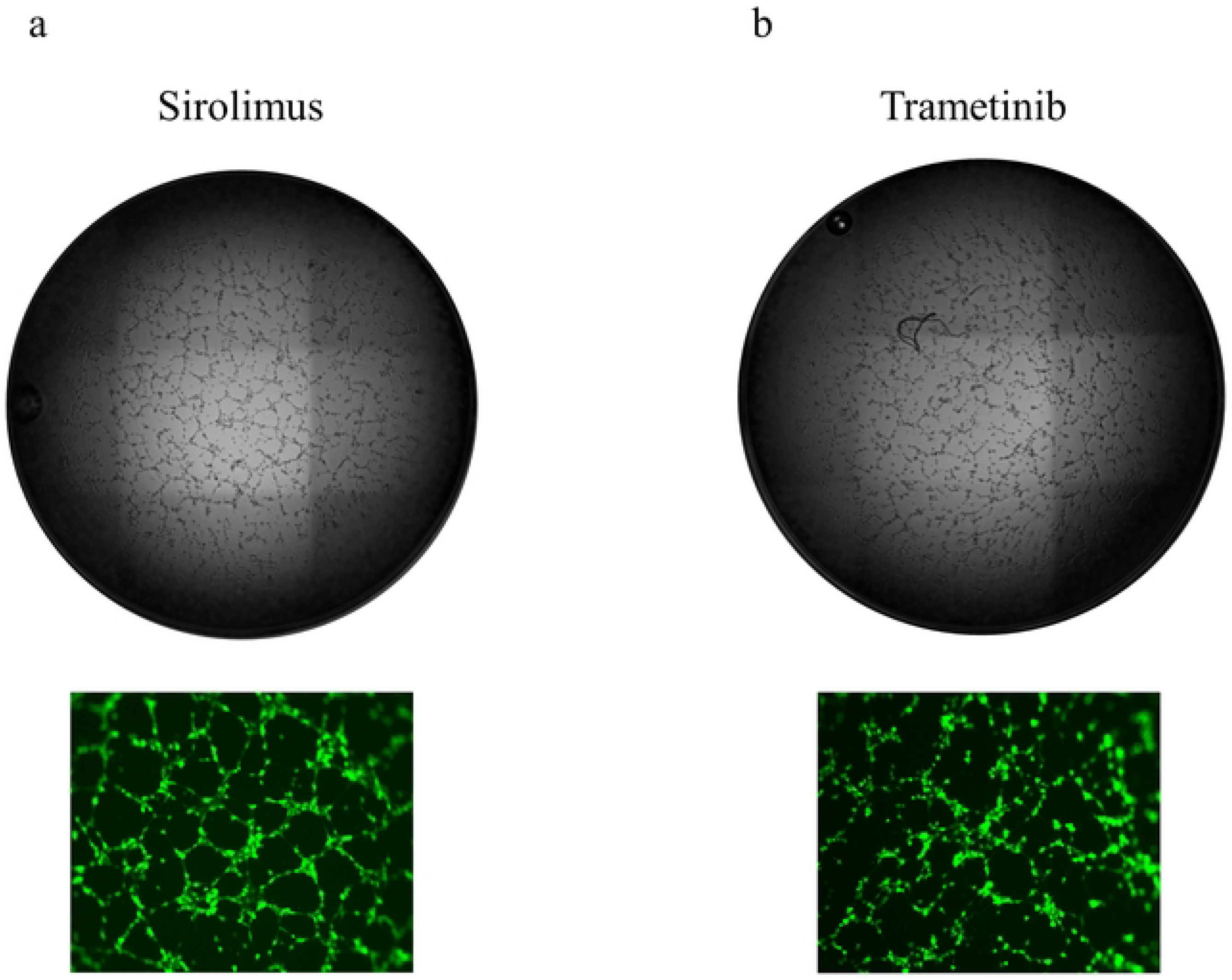
Tube formation assay of *NRAS*^WT^ and *NRAS^Q61R^* HDLECs. A: Ratios of *NRAS*^WT^, *NRAS^Q61R^*, and mixed HDLECs, and microscopy images. Cells were mixed at the indicated ratios and tube formation assays were performed. *NRAS*^WT^ HDLECs showed well-regulated lumen structures and tube formation, but *NRAS^Q61R^* HDLECs exhibited sheet-like proliferation and the formation of capillary-like structures was markedly less than that of *NRAS*^WT^ HDLECs. The higher the proportion of *NRAS^Q61R^* HDLECs (fluorescent cells), the greater the sheet-like proliferation and the fewer the capillary-like structures. 2B: In the stepwise examination of *NRAS^Q61R^* HDLEC mixtures, the covered area (occupied area by cellular components) (a), total nets (number of nets) (b), and tube length standard deviation (c) were analyzed by WinTube. Bars represent mean ± SD from triplicate wells. *p<0.05, **p<0.01, ***p<0.001, compared with 0%. 2C: Covered area (a), total nets (b), and tube length standard deviation (c) of *NRAS^Q61R^* HDLECs with or without treatment were analyzed by WinTube. Bars represent mean ± SD from triplicate wells. *p<0.05, **p<0.01, compared with *NRAS^Q61R^* HDLECs (no treatment). 2D: Microscopy images of tube formation of *NRAS^Q61R^* HDLECs treated with mTOR inhibitor sirolimus (30 ng/ml) (a) and MEK inhibitor trametinib (30 ng/ml) (b).

Time-lapse photography showed dynamic formational changes of the capillary-like structures of *NRAS*^Q61R^ HDLECs, and the cell migration was obviously different from that of *NRAS*^WT^ HDLECs (supplementary file. a, b). The greater the number of mutated cells, the more the formation of uniform and continuous loops was impaired in a concentration-dependent manner.

The covered areas occupying cellular components of 100% *NRAS*^Q61R^ HDLECs treated with sirolimus and trametinib were significantly smaller than those of the control (p=0.004 and p=0.004, respectively) (Fig. 2C-a). The number of nets of 100% *NRAS*^Q61R^ treated with sirolimus and trametinib was significantly higher than that of the control (p=0.015 and p=0.005, respectively) (Fig. 2C-b). The standard deviations of tube lengths of 100% *NRAS*^Q61R^ treated with sirolimus and trametinib were significantly shorter than those of the control (p=0.025 and p=0.025, respectively) (Fig. 2C-c). Sirolimus and trametinib treatment improved the poor tube-forming ability resulting from the *NRAS* mutation (Fig. 2D-a, b).

### Scratch assay

The scratch assay showed that *NRAS*^Q61R^ and *NRAS*^WT^ HDLECs had migrated into the void and completely closed the space. Migration of *NRAS*^Q61R^ HDLECs was more active than that of *NRAS*^WT^ and the space disappeared within 24 h (Fig. 3A). The space between wild type cells disappeared after 60 h. The space ratio of *NRAS*^Q61R^ at 0, 6, 12, 18, and 24 h decreased rapidly. However, that of *NRAS*^WT^ HDLECs showed a slow decrease (Fig. 3B).

**Figure 3.**
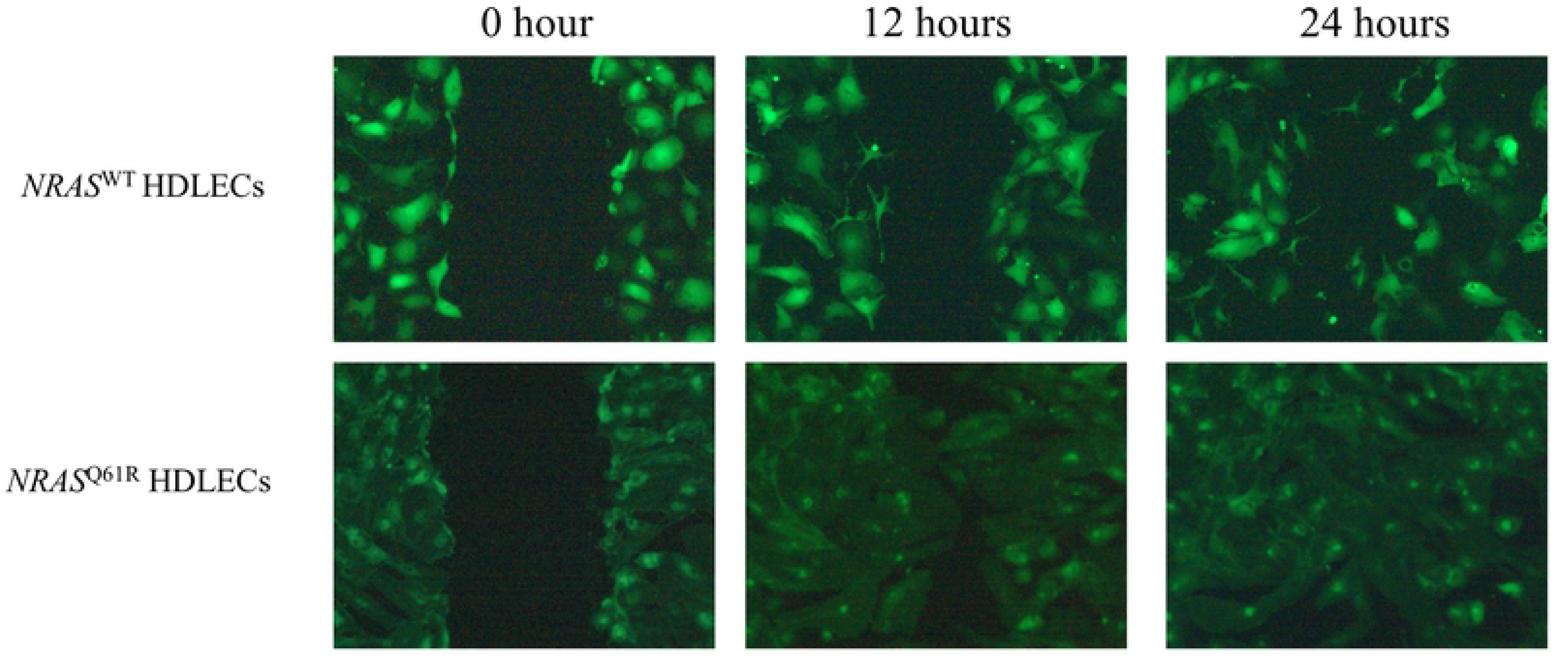
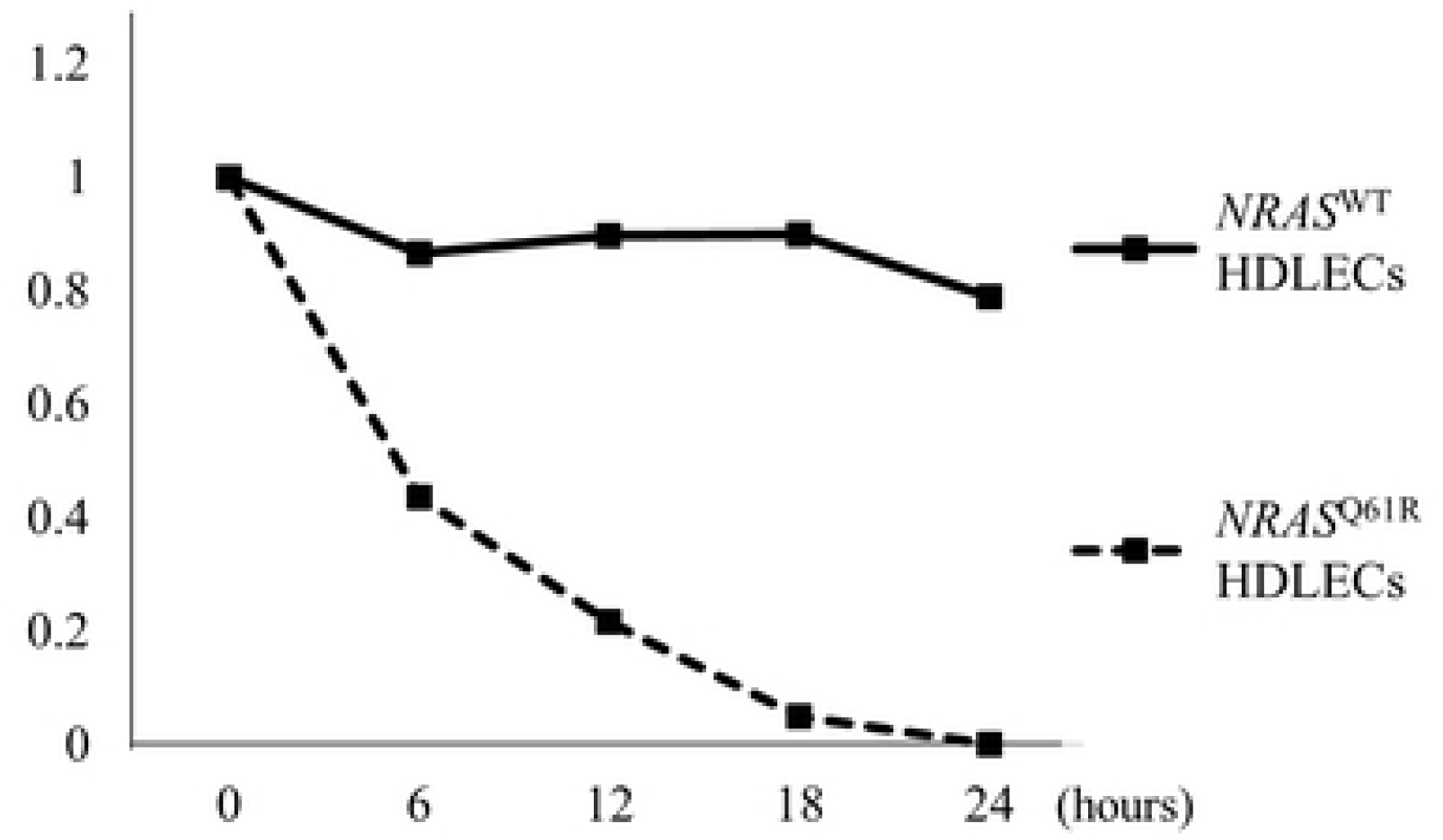
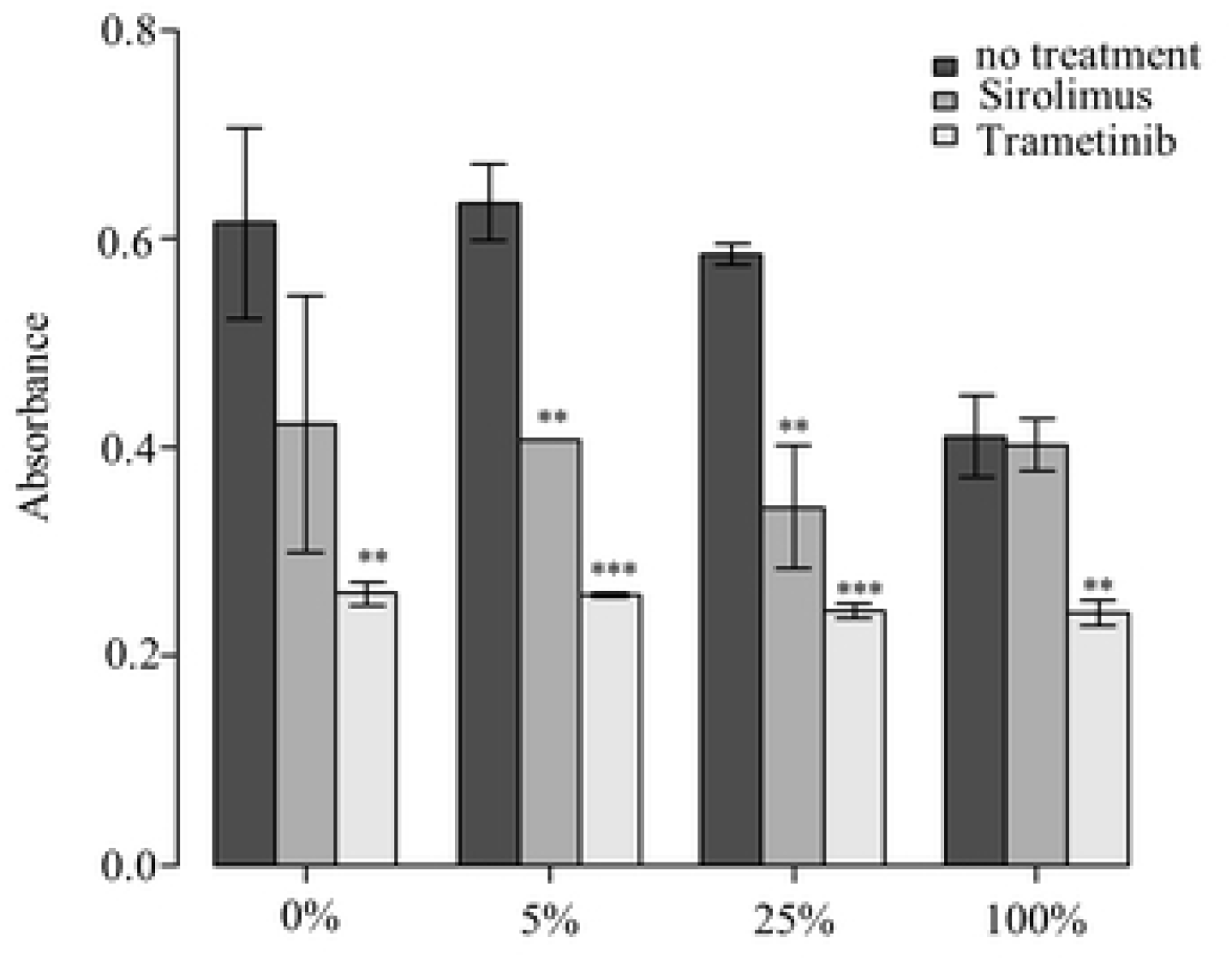
Scratch assay of *NRAS^Q61R^* and *NRAS*^WT^ HDLECs. 3A: Migration of *NRAS^Q61R^* HDLECs was more active than that of *NRAS*^WT^. The space ratio of *NRAS^Q61R^* HDLECs at 0, 6, 12, 18, and 24 h was decreased more rapidly than that of *NRAS*^WT^. 3B: Cell proliferation assay of *NRAS^Q61R^* and *NRAS*^WT^ HDLEC mixtures. 3C: Absorbance of *NRAS^Q61R^* HDLEC mixtures treated with sirolimus and trametinib. Bars represent mean ± SD from triplicate wells. *p<0.05, **p<0.01, ***p<0.001, compared with no treatment.

### Cell proliferation assay

The cell proliferation assay of *NRAS*^Q61R^ and *NRAS*^WT^ mixed HDLECs showed that the absorbance of 100% *NRAS*^Q61R^ was significantly lower than that of *NRAS*^WT^ (p=0.024) (Fig. 3C). There was no difference in absorbance between *NRAS*^WT^ and 5% and 25% *NRAS*^Q61R^. After sirolimus and trametinib treatment, the absorbance of *NRAS*^Q61R^ mixtures decreased significantly compared with that of cells with no treatment. There was no difference between the absorbance of 100% *NRAS*^Q61R^ after sirolimus treatment and that of no treatment, but the absorbance of 100% *NRAS*^Q61R^ after trametinib treatment was significantly lower than that of cells with no treatment (Fig. 3C).

### Analysis of signaling pathways in HDLECs by western blotting and multiplex protein assay

To determine the effects of *NRAS* mutation on the PI3K/AKT/mTOR and RAS/MAPK/ERK pathways, AKT, p-AKT, ERK1/2, and p-ERK1/2 expression was measured by western blot analysis. The expression levels of AKT in *NRAS*^Q61R^ were similar to those of *NRAS*^WT^ (Fig. 4A). However, the expression levels of p-AKT were slightly reduced in *NRAS*^Q61R^. ERK1/2 and p-ERK/1/2 levels in *NRAS*^Q61R^ were significantly upregulated.

**Figure 4.**
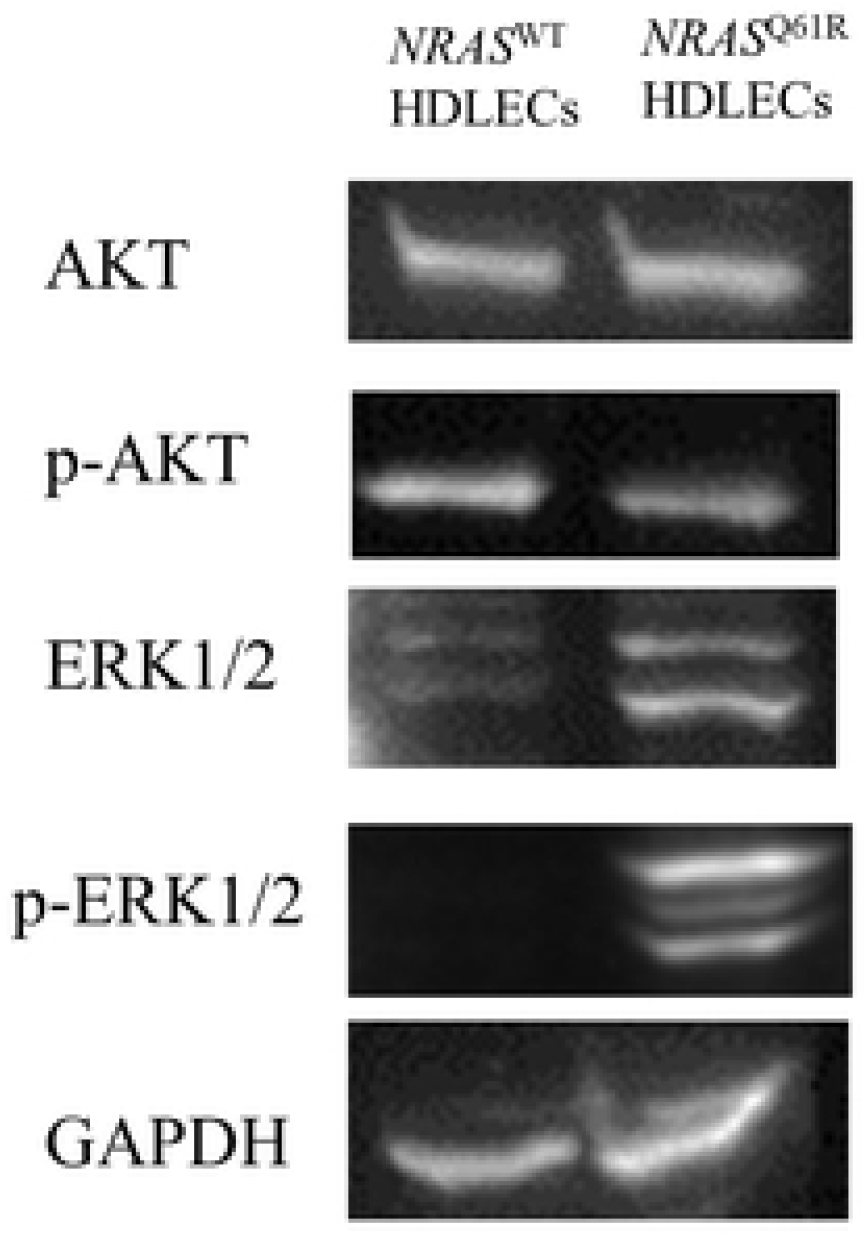
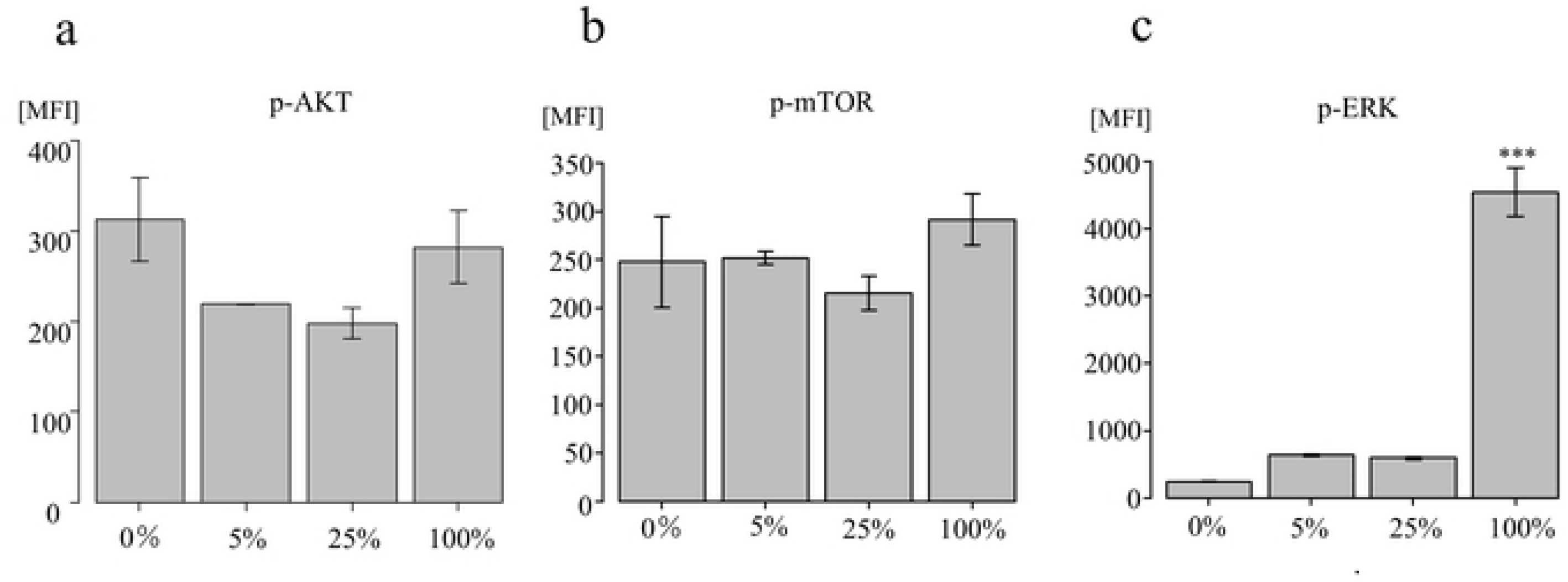
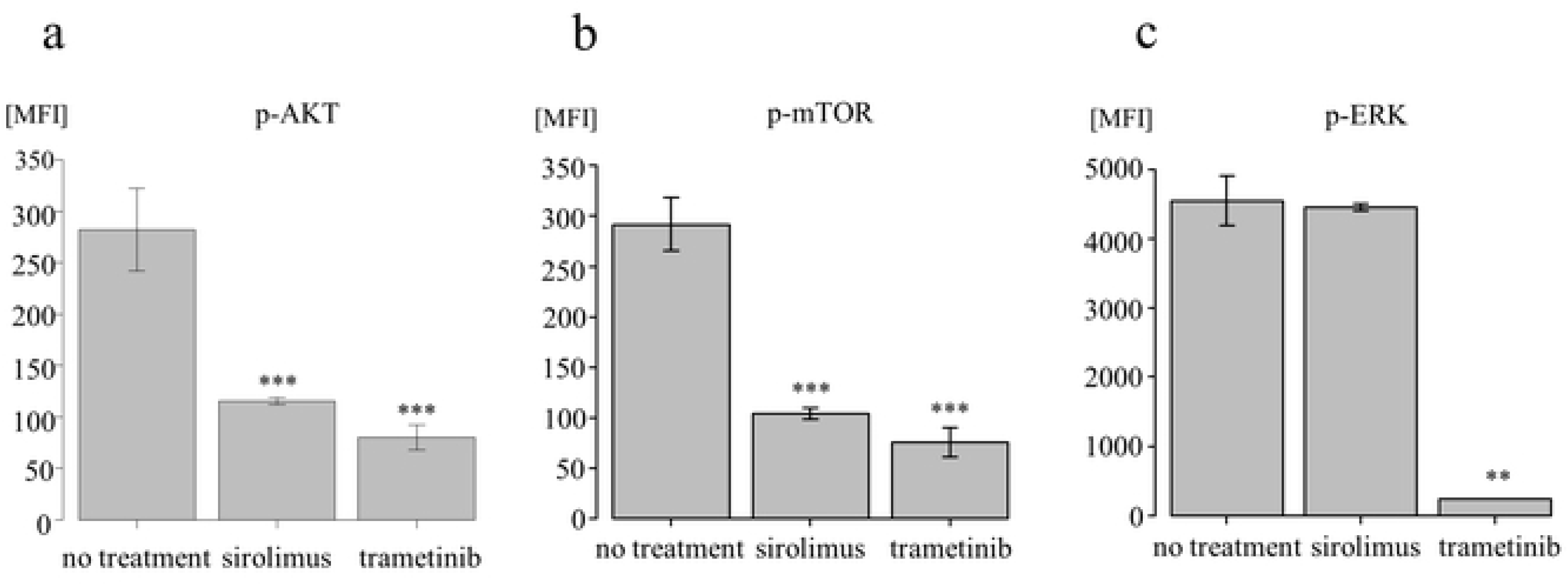
Analysis of signaling pathways in HDLECs by western blotting and multiplex protein assay. 4A: Western blot analysis of AKT, phospho-AKT (Ser473) (p-AKT [Ser473]), ERK1/2, and phospho-ERK1/2 (p-ERK1/2) expression in *NRAS^Q61R^* and *NRAS*^WT^ HDLECs 4B: Multiplex protein assay of IRS1 (a), AKT (b), p-mTOR (c), p70S6K (d), and p-ERK (e) in *NRAS^Q61R^* and *NRAS*^WT^ HDLEC mixtures. Bars represent mean ± SD from triplicate wells. *p<0.05, **p<0.01, ***p<0.001, compared with 0%. 4C: Multiplex protein assay of AKT (a), p-mTOR (b), and p-ERK(c) in *NRAS^Q61R^* HDLECs with or without treatment with sirolimus and trametinib. Bars represent mean ± SD from triplicate wells. *p<0.05, **p<0.01, ***p<0.001, compared with no treatment. After sirolimus and trametinib treatment, the absorbance of *NRAS*^Q61R^ HDLECs treated with sirolimus and trametinib decreased significantly compared with that of cells with no treatment.

Additionally, to quantitatively measure protein expression levels, we performed multiplex protein assay. The median fluorescence intensity (MFI) of IRS-1 decreased significantly even in small amounts of *NRAS*^Q61R^ HDLEC mixtures (p<0.001, p<0.001, and p<0.001, respectively) (Fig. 4B-a). The MFI of AKT, the major protein of the PIK3 pathway, was decreased in the cell group containing 5% and 25% *NRAS*^Q61R^ (p=0.021 and p=0.007, respectively), but there was no statistically significant difference between that of *NRAS*^WT^ and *NRAS*^Q61R^ (Fig.4B-b). The MFI of phosphorylated mTOR protein (p-mTOR) did not change depending on the ratio of *NRAS*^Q61R^ (Fig. 4B-c). The MFI of p70S6K, one of the proteins downstream of mTORC1, in *NRAS*^Q61R^ HDLEC mixtures decreased significantly compared with that of *NRAS*^WT^ (p<0.001, p<0.001, and p=0.002, respectively) (Fig. 4B-d). The MFI of phosphorylated ERK (p-ERK) was significantly increased in *NRAS*^Q61R^ (p<0.001) (Fig.4B-e). To assess the effects of sirolimus and trametinib treatment, we examined the expression of AKT, p-mTOR, and p-ERK. The MFI of AKT after sirolimus and trametinib treatment was significantly decreased compared with that of no treatment (p=0.001 and p<0.001, respectively) (Fig. 4C-a). The MFI of AKT after trametinib treatment was significantly decreased compared with that of no treatment (p<0.001 and p<0.001, respectively) (Fig. 4C-b). The MFI of p-ERK was significantly decreased after trametinib treatment (p=0.065), but was not decreased after sirolimus (Fig. 4C-c).

### Suspension array

Angiopoietin (ANG)-2, vascular endothelial growth factor (VEGF)-C, vascular endothelial growth factor receptor (VEGFR)-1, heparin binding-epidermal growth factor-like growth factor (HB-EGF), VEGFR-2, interleukin (IL)-8, and europilin-1 levels in the medium of both *NRAS*^WT^ and *NRAS*^Q61R^ HDLECs were significantly higher than those of the controls (Fig. 5a–d; Sup Fig. 1a–c). ANG-2, VEGFR-1, HB-EGF, and neuroplin-1 expression in *NRAS*^Q61R^ was significantly higher than in *NRAS*^WT^ HDLECs (Fig. 5a, c, d; Sup Fig. 1c). VEGF-C and VEGFR-2 levels in *NRAS^Q61R^* were significantly lower than in *NRAS*^WT^ (Fig.5b; Supp Fig.1a). There was no significant difference in IL-8 levels between *NRAS*^WT^ and *NRAS^Q61R^* (Supp Fig.1b). G-CSF, SCF/c-kit, and TIE2 expression was similar in *NRAS*^WT^ and the control, and significantly higher in *NRAS^Q61R^* (Fig. 5e, Sup. 1d, e). The VEGF-A levels of *NRAS*^WT^ and *NRAS^Q61R^* were lower than those of the control (Supp Fig.1f).

**Figure 5.**
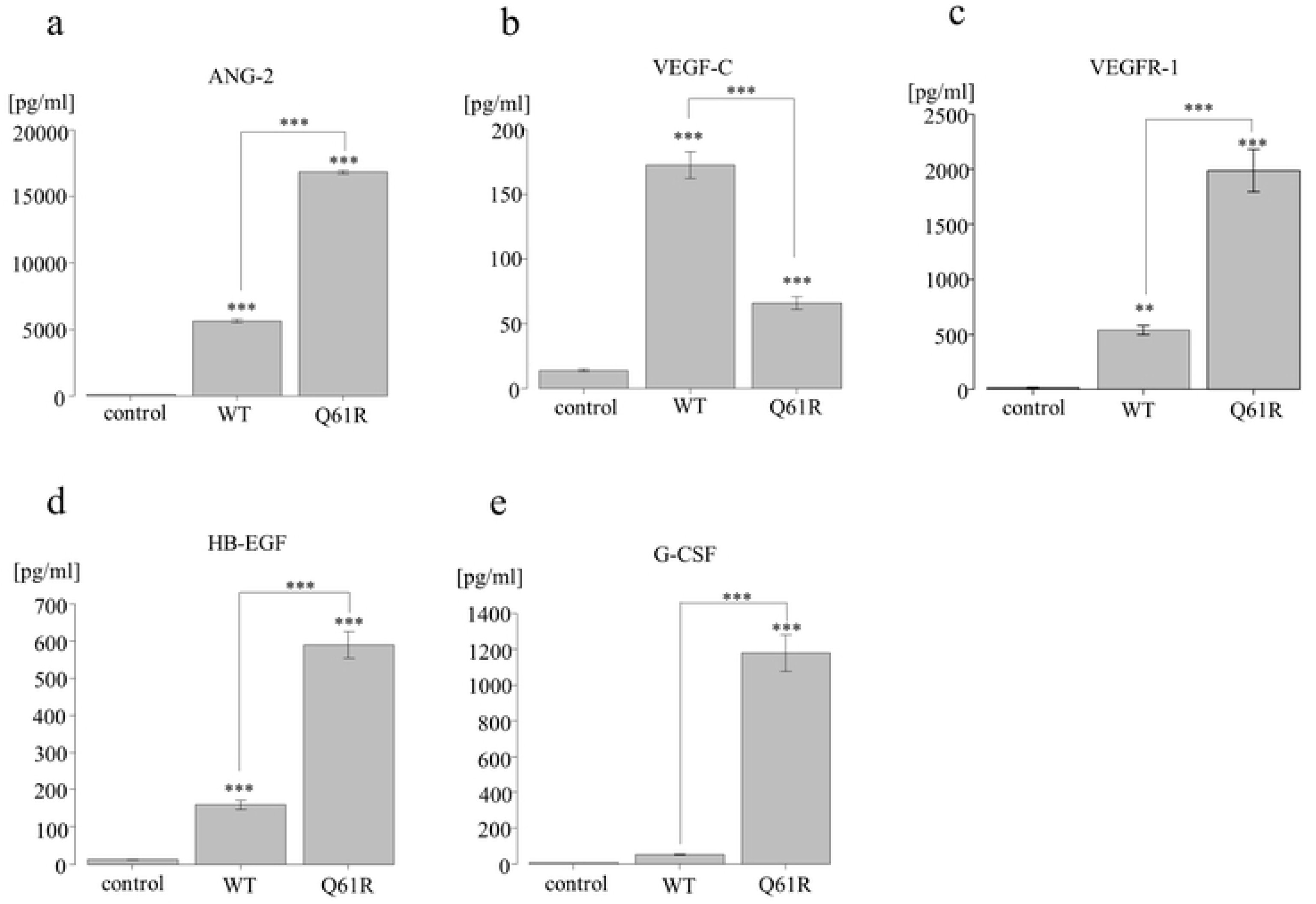
Suspension array of *NRAS^Q61R^* and *NRAS*^WT^ HDLECs. Angiopoietin-2 (ANG-2) (a), vascular endothelial growth factor (VEGF)-C (b), vascular endothelial growth factor receptor (VEGFR)-1 (c), heparin binding-epidermal growth factor-like growth factor (HB-EGF) (d), and granulocyte-colony stimulating factor (G-CSF) (e) in *NRAS^Q61R^* and *NRAS*^WT^ HDLECs. Bars represent mean ± SD from triplicate wells. *p<0.05, **p<0.01, ***p<0.001, compared with control. *NRAS^Q61R^* and *NRAS*^WT^ HDLECs were also compared.

### Metabolomics of HDLECs

The number of confluent cells in the 10-cm dishes was 1.3 × 10^6^ cells for *NRAS*^WT^ HDLECs and 0.4 × 10^6^ cells for *NRAS*^Q61R^ HDLECs. As mentioned above, there is a large difference in the size of the two cell lines, and therefore confluent cells in the 10-cm dishes were analyzed. The total amount of amino acids were almost the same in the two lines, with 213587.0 pmol/1.3 × 10^6^ cells in *NRAS*^WT^ and 234899.6/0.4 × 10^6^ cells in *NRAS*^Q61R^ HDLECs.

Among the tricarboxylic acid (TCA) cycle, citric acid, cis-aconitic acid, isocitric acid, and α-ketoglutaric acid showed significantly higher values in *NRAS*^Q61R^ than in *NRAS*^WT^ HDLECs (p=0.007, p=0.016, p=0.016, and p=0.006, respectively) (Fig.6). There was no significant difference in succinic acid and fumaric acid. Malic acid was significantly higher in *NRAS*^WT^ (p=0.037). The 2-hydroxyglutaric acid synthesized from α-ketoglutaric acid was also not significantly different between *NRAS*^Q61R^ and *NRAS*^WT^.

**Figure 6.**
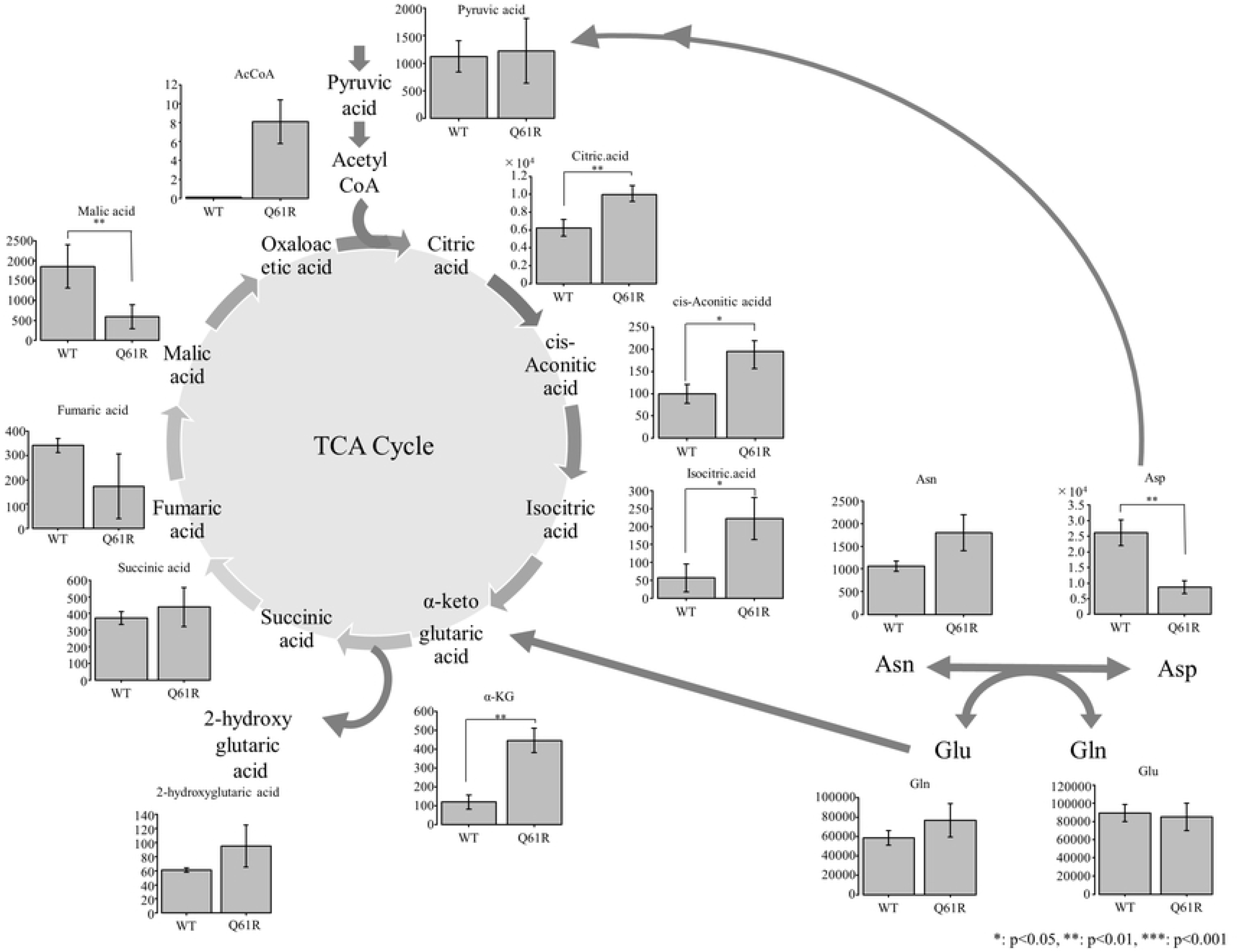
Metabolome analysis of the TCA cycle in *NRAS^Q61R^* and *NRAS*^WT^ HDLECs. Bars represent mean ± SD from triplicate wells. *p<0.05, **p<0.01, ***p<0.001

Among other amino acids, aspartic acid in *NRAS*^WT^ HDLECs was significantly higher than in *NRAS*^Q61R^ (p=0.008) (Fig.5). Asparagine and glutamic acid (Glu) in *NRAS*^Q61R^ were slightly higher than in *NRAS*^WT^, but there was no significant difference. Glutamine in both cell lines did not show a significant difference.

## Discussion

We herein report that *NRAS*^Q61R^ HDLECs had a higher migration ability and a lower cell proliferation ability than *NRAS*^WT^. Tube formation became poorer with increasing proportions of *NRAS*^Q61R^ HDLECs. Activation of the PIK3/AKT/mTOR pathway in *NRAS*^Q61R^ HDLECs was suppressed and poor tube formation was improved by mTOR inhibitor treatment. These results indicate the theoretical *in vitro* circumstances induced by *NRAS*^Q61R^ mutated cells in the affected lesions of KLA patients.

The pathological findings of affected lesions in KLA patients showed morphological abnormalities of lymphatic cells and clusters of spindle cells. Although the very low frequency NRAS p.Q61R variant (1%–28%) was detected in the samples of patients with KLA in previous studies^4^, it is unknown which type of cells have this variant. In our study, we transfected *NRAS*^Q61R^ vectors into HDLECs, which showed abnormal tube formation and proliferated in sheet form. In the mixed cells group, the higher the proportion of mutated cells, the larger the sheet-like area. Abnormal lymphatic tissue proliferation in KLA patients may also be caused by partial loss of normal lymphatic construction.

A previous study examined the characteristics of patient-derived cells from two KLA patients, one of whom had the same *NRAS*^Q61R^ mutation^7^. Tube formation assay showed that KLA cells had significantly fewer tubes than normal HDLECs. Their tubes proliferated in sheet form, similar to our *NRAS*^Q61R^ HDLECs^7^. Although it remains unclear as to why a small amount of mutated HDLECs can affect the progress of lesions in actual KLA patients, our results may show a hypothetical circumstance using mixed *NRAS*^Q61R^ HDLECs.

Recent studies indicated that the PIK3/AKT/mTOR and RAS/MAPK/ERK pathways are important in the pathogenesis of KLA^4, 5^. Western blotting revealed that the levels of p-AKT in *NRAS*^Q61R^ HDLECs were downregulated, but those of ERK1/2 and p-ERK1/2 were upregulated. In a multiplex protein assay, IRS-1 and p70S6K levels in *NRAS*^Q61R^ HDLECs were downregulated, but those of p-ERK1/2 were upregulated. These results showed that inactivation of the PIK3/AKT/mTOR pathway and hyperactivation of the RAS/MAPK/ERK pathway were caused by the activating *NRAS*^Q61R^ mutation. A recent study of human endothelial progenitor cells expressing *NRAS*^Q61R^ mutation also showed the same effects on these pathways^16^. However, established cell lines from three patients demonstrated various results including downregulation of p-ERK or upregulation of p-AKT^7, 17^. Although it is challenging to predict whether these pathways are activated or inactivated in affected lesions, these established cell lines may contain not only the mutated cells but also non-mutated cells that consist of pathological structures and have abnormal functions in the affected lesions. It is not yet clear how mutated cells and non-mutated cells correlate with each other.

Sirolimus is a first-line drug for refractory CLAs, and trametinib is an alternative drug candidate^18, 19^. In previous studies, these drugs inhibited the proliferation of patient-derived cells^7, 16^. Our studies showed that tube formation and cell proliferation of *NRAS*^Q61R^ HDLECs improved after both sirolimus and trametinib treatment. Trametinib could inhibit the PIK3/AKT/mTOR pathway because it is closely related to the RAS pathway^20^. However, the proliferation of 100% *NRAS*^Q61R^ was inhibited only by trametinib treatment, and not by sirolimus. In a multiplex protein assay, the MFI of AKT and p-mTOR following sirolimus treatment was significantly decreased compared with that of non-treated cells, but that of p-ERK was decreased only after trametinib treatment. Similarly, Boscolo et al. reported that a MEK inhibitor could improve measurement of the cellular shape of *NRAS*^Q61R^ mutant human endothelial cells, whereas mTOR inhibitor could not^16^. It is unclear why mTOR and MEK inhibitors can improve the tube formation and proliferation of patient-derived cells and mixed *NRAS*^Q61R^ HDLECs, and why only MEK inhibitor can prevent the proliferation of 100% *NRAS*^Q61R^ HDLECs and cell circularity.

ANG-2 and VEGF-C are pivotal factors related to lymphangiogenesis^21^. ANG-2, which binds to TIE2, is known to cause maturation and stabilization after remodeling of LECs^21–23^. VEGF-C, which binds to VEGFR-3, has an important role in vasculogenesis, sprouting, migration, and proliferation of lymphatic cells^21^. In our study, the levels of these proteins in the medium of both *NRAS*^WT^ and *NRAS*^Q61R^ HDLECs were higher than those of the controls. ANG-2 levels in the medium of *NRAS*^Q61R^ were significantly higher, but VEGF-C levels were lower than that of *NRAS*^WT^. Tube formation assay showed that *NRAS*^Q61R^ HDLECs proliferated excessively and formed tubes poorly. Therefore, we considered that hypersecretion of ANG-2 from *NRAS*^Q61R^ HDLECs may cause decreasing VEGFR-3 levels and might be related to inhibition of the maturation of peripheral LECs. Furthermore, in previous reports of the cytokine profiles of KLA patients, ANG-2 levels were elevated in the patients’ samples, suggesting it could be an important biomarker of KLA^24, 25^. Although the association with these cytokines and diseases is unknown, ANG-2 secretion and activation of these signaling cascades might be associated with the pathogenesis of KLA.

Growth factors such as G-CSF and HB-EGF in *NRAS*^Q61R^ HDLECs were also significantly higher than those in *NRAS*^WT^. In cancer cells, it is considered that growth factors and cytokines produced by cells within the tumor microenvironment might activate G-CSF in tumor and stromal cells because activation of the RAS/RAF/MEK pathway resulted in enhanced G-CSF expression^26^. HB-EGF is also thought to play a pivotal role in cell proliferation and differentiation. In particular, soluble HB-EGF is a potent promoter of cell adhesion, cell motility, and angiogenesis^27^. However, although the relationship between these growth factors and the pathogenesis of vascular anomalies has not been elucidated, growth factors secreted by some *NRAS*^Q61R^ HDLECs in affected lesions might have influenced not only the proliferation of mutated cells but also abnormalities of peripheral normal LECs.

Many reports have shown that RAS is associated with metabolic changes in various cancer cell types^28–30^ and mTOR is known to be involved in glucose and amino acid metabolism^31^. We comprehensively analyzed metabolites and compared *NRAS*^WT^ HDLECs with *NRAS*^Q61R^. In the first half of the TCA cycle, which involves citric acid, cis-aconitic acid, isocitric acid, and α-ketoglutaric acid, the metabolite levels were significantly higher in *NRAS*^Q61R^. α-Ketoglutaric acid is produced from isocitric acid in the TCA cycle and also from Glu. Glu that is produced from Gln becomes α-ketoglutaric acid and enters the TCA cycle. Glutaminolysis is a metabolic pathway that extracts energy from glutamine, which is unique to cancer cells. It has been reported that glutaminolysis is caused by activation of *KRAS* in various cancers^30^. Thus, glutaminolysis is considered one of the major energy supply pathways in RAS-driven cancer^28^. Although it is not clear how glutaminolysis is associated with the RAS/MAPK/ERK pathway, our metabolome analysis demonstrated that *NRAS* mutation can lead to changes in cellular energy metabolism.

In summary, *NRAS*^Q61R^ HDLECs showed poor tube formation, low cell proliferation, and high migration ability, and an increasing ratio of mutated cells led to poorer tube formation. mTOR and MEK inhibitors can improve tube formation and decrease cell proliferation. Mutations of the *RAS* genes of LECs drive incessant activation of critical pathways, which in turn dysregulates the function of cytosolic and nuclear signaling effector molecules such as cytokines and metabolic regulatory proteins. This might cause abnormalities in the tissue function and differentiation of the affected lesions of CLAs.

## Acknowledgements

We thank the Department of Pediatrics at Gifu University for their contributions. We also thank Ms. Asuka Ogawa of Gifu University for providing technical assistance. The present study was supported in part by a Practical Research Project for Rare/Intractable Diseases (JP22ek0109515) from Japan’s Agency for Medical Research and Development, AMED. Finally, we thank H. Nikki March, PhD, from Edanz (https://jp.edanz.com/ac) for editing a draft of this manuscript.

---

**Supplemental Figure 1.**
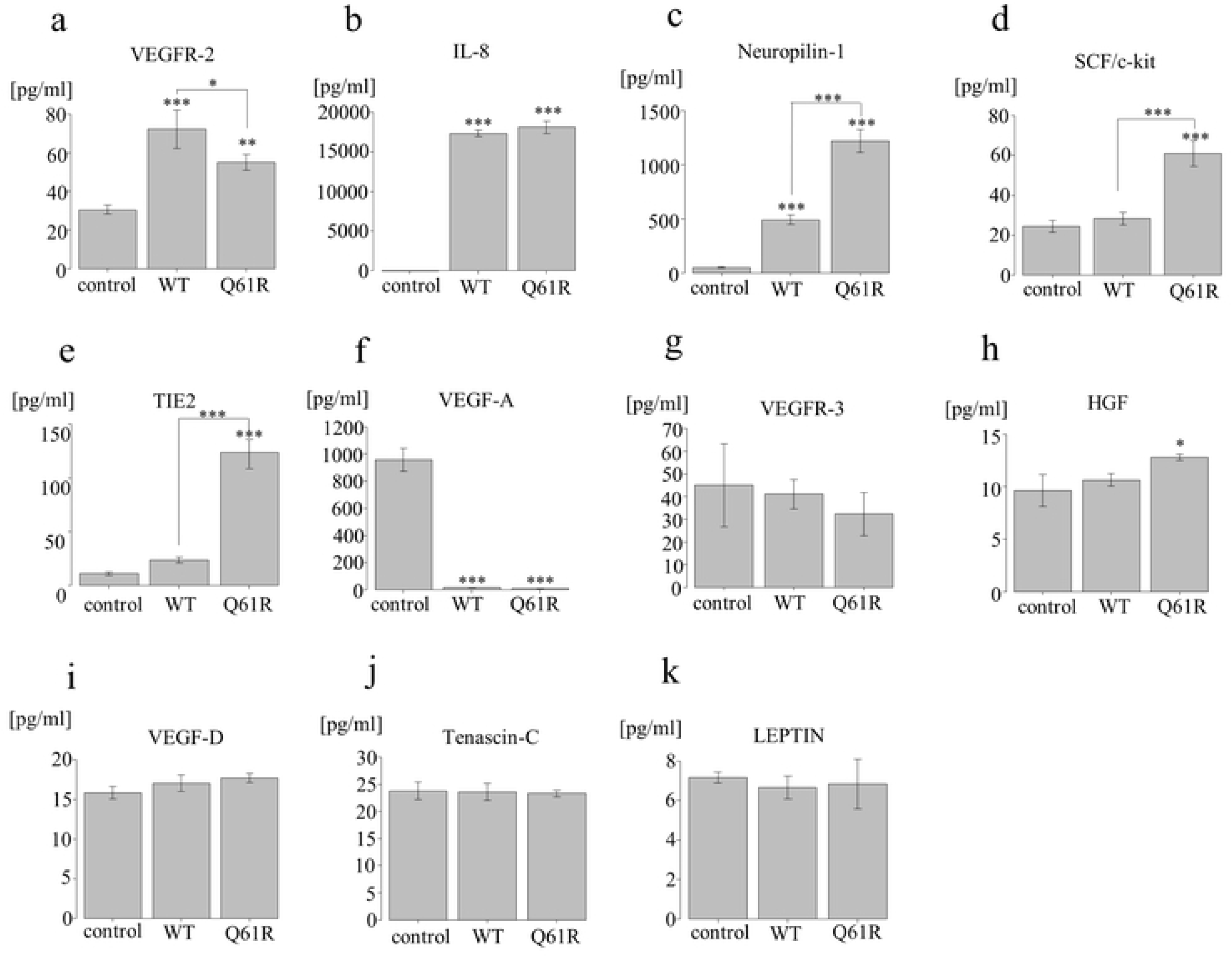
Suspension array of *NRAS^Q61R^* and *NRAS*^WT^ HDLECs. Vascular endothelial growth factor receptor (VEGFR)-2 (a), interleukin (IL)-8 (b), neuroplin-1 (c), stem cell factor (SCF)/c-kit (d), TIE2 (e), vascular endothelial growth factor (VEGF)-A (f), VEGFR-3 (g), hepatocyte growth factor (HGF) (h), VEGF-D (i), tenascin-C (j), and leptin (k). Bars represent mean ± SD from triplicate wells. *p<0.05, **p<0.01, ***p<0.001, compared with control. *NRAS^Q61R^* and *NRAS*^WT^ HDLECs were also compared.

## Supplementary files

Time-lapse photography showing dynamic formational changes of *NRAS*^WT^ HDLECs (a) and *NRAS*^Q61R^ HDLECs (b). *NRAS*^Q61R^ HDLECs showed impaired formation of uniform and continuous loops.

